# Neuronally Produced Betaine Acts via a Novel Ligand Gated Ion Channel to Control Behavioural States

**DOI:** 10.1101/2021.10.29.466399

**Authors:** I Hardege, J Morud, J Yu, TS Wilson, FC Schroeder, WR Schafer

## Abstract

Trimethyl glycine, or betaine, is an amino acid derivative found in diverse organisms, from bacteria to plants and animals. It can function as an osmolyte to protect cells against osmotic stress, and building evidence suggests betaine may also play important functional roles in the nervous system. However, despite growing interest in betaine’s roles in the nervous system, few molecular mechanisms have been elucidated. Here we identify the expression of betaine synthesis pathway genes in the nervous system of the nematode worm, *C. elegans*. We show that betaine, produced in a single pair of interneurons, the RIMs, can control complex behavioural states. Moreover, we also identify and characterise a new betaine-gated inhibitory ligand gated ion channel, LGC-41, which is required for betaine related behavioural changes. Intriguingly we observed expression of LGC-41 in punctate structures across several sensory and interneurons, including those synaptically connected to the RIMs. Our data presents a neuronal molecular mechanism for the action of betaine, via a specific receptor, in the control of complex behaviour within the nervous system of *C. elegans*. This may suggest a much broader role for betaine in the regulation of animal nervous systems than previously recognised.

## Main

The amino acid derivative betaine is best characterised for its role in osmoregulation. In mammals, betaine is an important osmolyte in the kidney medulla, where its transport is essential for adaptation to hypertonic stress^1^. Even in single celled organisms, betaine acts as an osmolyte, with many bacterial species expressing betaine transporters which are critical for maintaining cellular osmolarity^2^. In the mammalian liver betaine acts as a methyl donor during the transmethylation of homocysteine to methionine^3^. However, the presence of the selective betaine and GABA transporter, BGT1^4^, which has relatively low affinity for GABA compared to other GABA specific transporters^5^, has led to a growing interest in the role of betaine in regulating the nervous system. A building body of evidence suggests betaine may indeed have specific roles in the nervous system, for example, dietary supplementation with betaine has been shown to improve cognitive performance in humans as well as improving memory in rodents^6^. In addition, betaine has been shown to have anticonvulsant properties, such as inhibiting pharmacologically-induced seizures in the rat^7,8^. Moreover, inhibitors of BGT1, expressed in neuronal dendrites, have shown promise as potential anti-epileptic drugs^8^. Although these effects could be mediated indirectly, for example through effects on local concentrations of GABA or on neuronal metabolism, the effects are also consistent with a direct action of betaine on neurons. However, a molecular target for betaine in mammalian neurons has not yet been identified.

Interestingly, the nematode *C. elegans* expresses a homolog of BGT1, *snf-3*, which encodes a betaine-selective transporter, without measurable activity toward GABA^9^. In addition *C. elegans* also expresses a muscular ligand gated ion channel, *acr-23*, that is activated by betaine^9,10^. However, the relevant source of betaine in nematodes, and whether it has a functional role in the nematode nervous system, has remained unclear. In addition, it is not known whether betaine is synthesised and released in an activity-dependent manner from nematode neurons to modulate neural circuits and behaviour.

### LGC-41 is an Inhibitory Betaine Receptor Expressed Broadly in The Nervous System

In comparison to mammals, the *C. elegans* genome contains an expanded number of ligand gated ion channel (LGIC) genes^11^. By performing an unbiased ligand screen of orphan channels, where LGICs were expressed in *Xenopus* oocytes and screened for induction of current in the response to a panel of potential ligands, we identified a new betaine-gated ion channel, LGC-41 (Fig. 1A-B). LGC-41 belongs to a diverse gene family of LGICs in *C. elegans*, whose other members include choline, acetylcholine, and monoamine-gated chloride channels^12^ (Fig. 1A). Of the panel of neurotransmitters we tested, LGC-41 was specifically gated by betaine (Fig. 1A-B), with an EC_50_ of 211 *µ*M (Fig. 1C), we saw no activity in the presence of acetylcholine, choline, or any of the other tested neurotransmitters (Fig. 1B). Using ion substitution experiments we identified LGC-41 as an inhibitory chloride channel, in line with the rest of this subfamily (Fig. 1D). LGC-41 was not effectively blocked by any cholinergic blockers at low concentrations (Fig. 1E). Only strychnine, traditionally considered a glycine receptor antagonist, was able to block LGC-41 with an IC_50_ of 973 *µ*M (Fig. 1E). Interestingly, LGC-41 could be blocked by the anion pore blocker picrotoxin with an IC_50_ of 144 *µ*M (Fig. 1E). When exposed to multiple applications of betaine we saw no reduction in peak current size between pulses of 10s, 30s or 60s intervals, suggesting that the receptor is accessible for reactivation after 10s (Fig. 1F).

**Figure 1.**
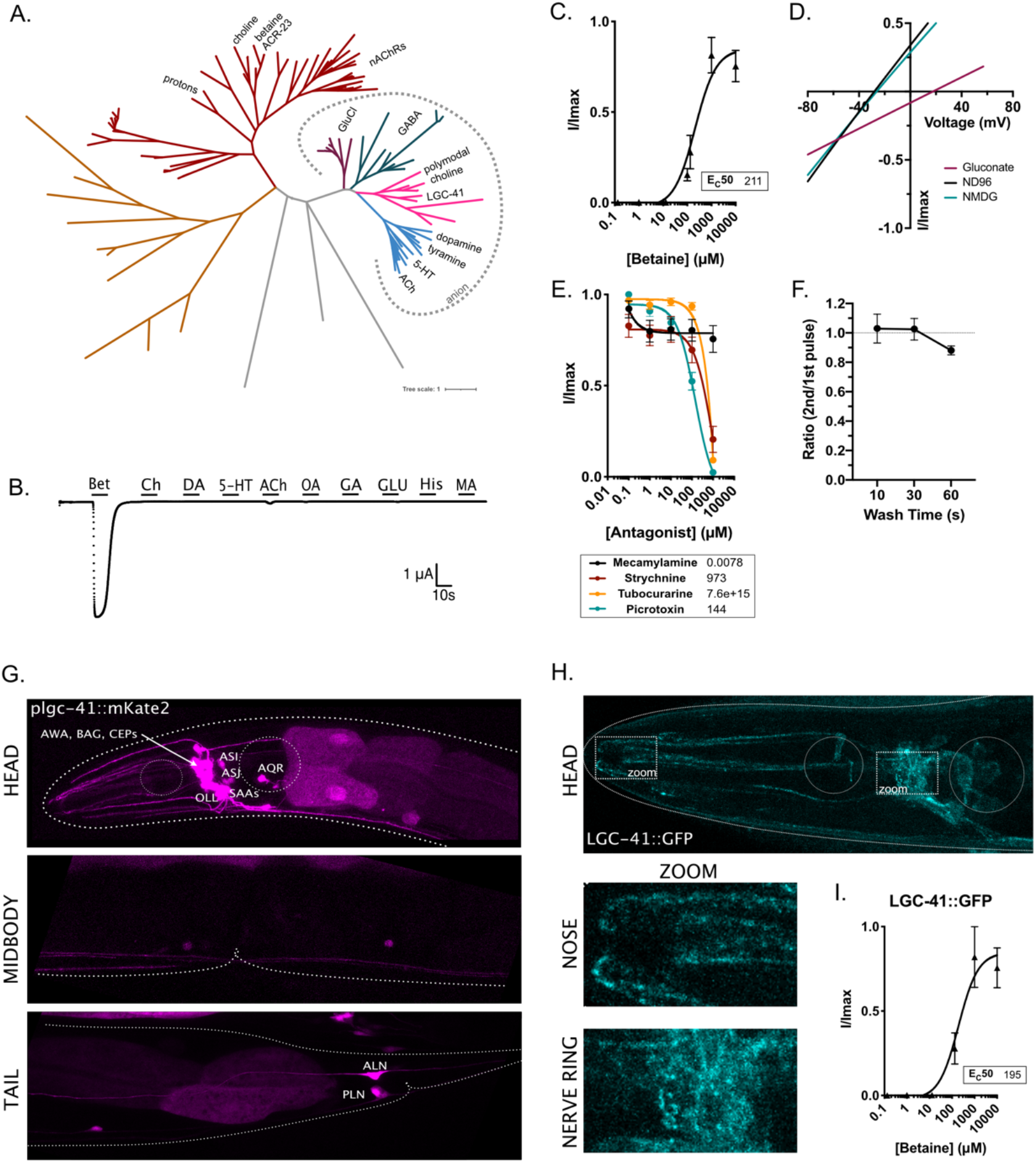
LGC-41 forms an inhibitory betaine-gated channel. (**A**) Phylogenetic tree showing evolutionary relationship between *C. elegans* LGICs, branch colors show subfamilies as follows: nAChRs (red), uncharacterized (orange and grey), GluCl (purple), GABA-gated anion and cation channels (green), diverse channels including LGC-41 (pink), ACh-gated and monoamine gated channels (blue). Figure made with iTOL using an un-rooted tree with 1 iteration of the equal-daylight algorithm (**B**) Continuous current trace for an oocyte expressing LGC-41 held at -60mV and perfused with a panel of ligands. (**C**) Dose response curve for oocytes expressing LGC-41 exposed to increasing concentrations of betaine, error bars represent SEM of at least 5 oocytes. Insert shows EC_50_ in *µ*M calculated with 3 parameter Hill slope. (**D**) Representative plot of current-voltage relationship of betaine induced current in oocytes expressing LGC-41 in ND96 (Na^+^ & Cl^-^ present), Na Gluconate (low Cl^-^) and NMDG (no Na^+^). (**E**) Antagonist dose response curves in oocytes expressing LGC-41 in the presence of 500 *µ*M betaine and increasing concentrations of each antagonist, error bars represent SEM of at least 7 oocytes. Insert shows IC_50_ in *µ*M calculated with 3 parameter Hill slope. (**F**) Ratio of betaine induced current in oocytes expressing LGC-41 exposed to multiple pulses of 500 *µ*M betaine with 10, 30 and 60s wash time between pulses. Error bars represent SEM of at least 6 oocytes. (**G**) Fluorescent reporter of mKate2 driven under the promoter sequence of *lgc-41*. (**H**) Images of LGC-41::GFP tagged endogenously with CRISPR/Cas9 in the M3/4 loop, arrows point to zoomed areas which show localisation in the nose and nerve ring. (**I)** Dose response curve for oocytes expressing LGC-41::GFP exposed to increasing concentrations of betaine, error bars represent SEM of at least 4 oocytes. Insert shows EC_50_ in *µ*M calculated with 3 parameter Hill slope.

To determine the function of LGC-41 within the nervous system we characterised the expression of the channel using fluorescent reporters under the control of the *lgc-41* promoter. This revealed broad expression of *lgc-41* in several neuron classes including the sensory neurons AWA, BAG, CEP, AQR, OLL, ASJ and ASI (Fig. 1G). Many of these neurons are known to be involved in the regulation of the behavioural response to changes in the environment; AQR and BAG regulate oxygen sensation and social feeding^13^, the CEPs induce behavioural changes in response to food sensation^14^ and ASI is involved in integrating multiple sensory inputs to regulate feeding and food search behaviours^15^. LGC-41 was also found to be expressed in SAA interneurons, which are important in controlling steering and navigation^16^ (Fig. 1G). Using a functional GFP tagged LGC-41 (Fig 1I), generated by CRISPR insertion, we show endogenous localisation of LGC-41 in both clear punctate structures and diffuse signal, being present both in the synaptically dense nerve ring region as well as in what appears to be sensory nerve endings in the nose of the worm (Fig. 1H). This localisation suggests a possible neuronal function for LGC-41 and its ligand, betaine, in neuronal communication or chemosensation.

### LGC-41 Promotes Global Search Behaviour

To examine the role of LGC-41 in the *C. elegans* nervous system we characterised the behaviour of *lgc-41* null animals. Strikingly, we observed that, despite performing typical feeding behaviour in the presence of food, *lgc-41* mutant animals remained in the centre of a plate in the absence of food in large numbers, which is in stark contrast to the typical dispersal and search behaviours initiated by wildtype worms in the absence of food. *C. elegans* transition between two distinct search behaviours in the absence of food, the first, local search or area restricted search, lasting approximately 15 min consists of frequent omega turns and short reversals. The second, global search or dispersal, is characterised by long forward runs^15,17^. Reasoning that the lack of dispersal from the centre of the plate indicated a change in this search state transition we systemically examined whether *lgc-41* mutant worms were able to perform the switch between the two search behaviours, local and global search. We placed a high density of washed worms in the centre of an empty plate containing no bacteria and observed whether they dispersed to explore the area of the plate after 1 hour (Fig 2A). Indeed, we found that *lgc-41* mutants were significantly less likely to disperse across the plate than N2 (Fig 2B-C).

**Figure 2.**
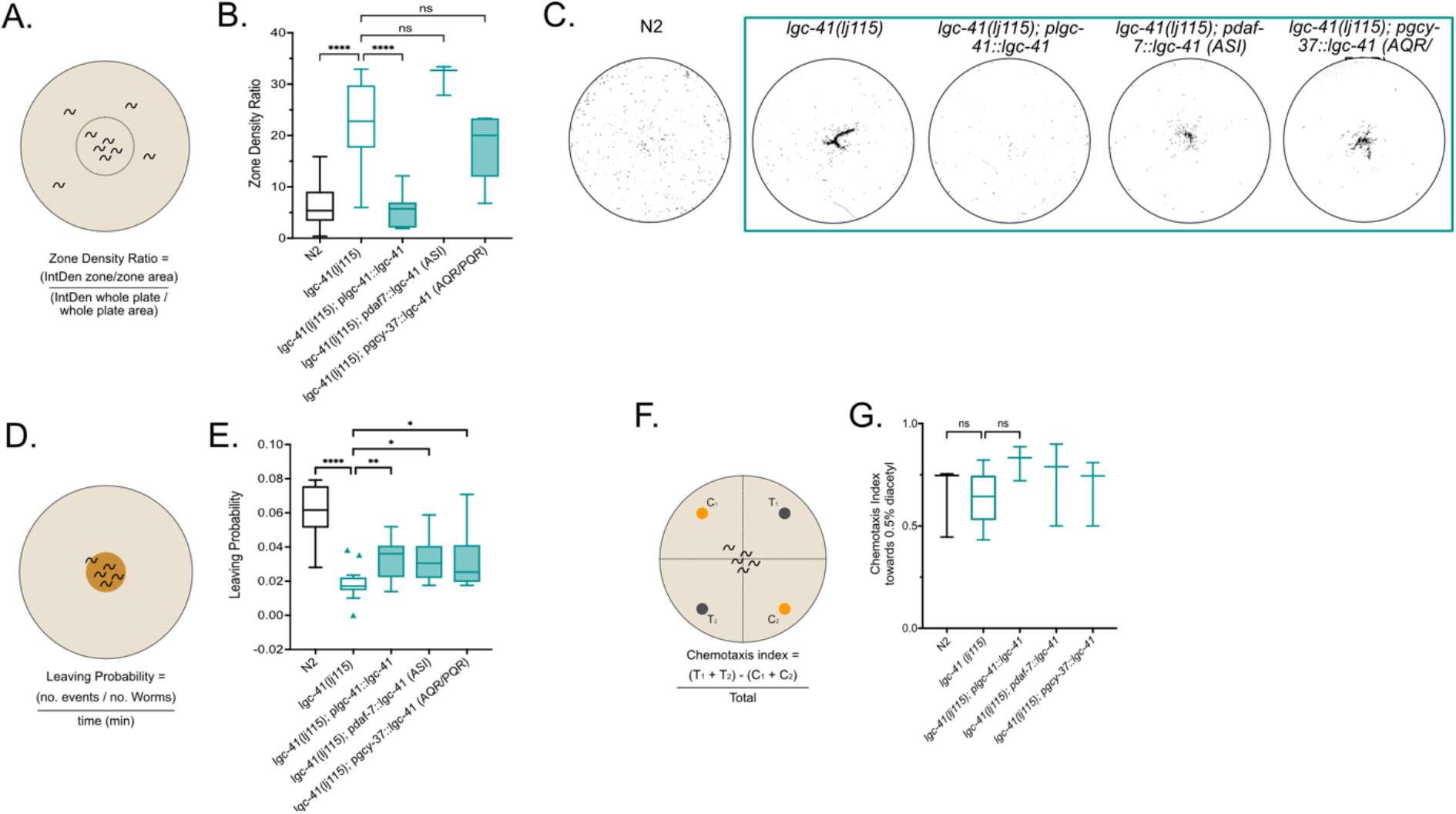
LGC-41 is required for global search and food leaving behaviours. Behavioural responses of N2, *lgc-41(lj115), lgc-41(lj115); plgc-41::lgc-41, lgc-41(lj115); pdaf-7(ASI)::lgc-41, lgc-41(lj115); pgcy-37(AQR/PQR)::lgc-41*. (**A**) Schematic representation of the experimental design and (**B**) box plot of dispersal assay, in which central dispersal index is calculated and plotted. N=3-29 plates per genotype. (**C**) Representative images of threshold filtered plates used to calculated dispersal index. (**D**) Schematic representation of the experimental design and (**E**) box plot of food leaving probability 6h after being placed on the food patch. N=16-21 plates per genotype. (**F**) Schematic representation of experiment (**G**) Chemotaxis index towards 0.5% diacetyl. N= 3-5 plates per genotype. (**B, C, E, G**) Tukey’s blot plots and one-way ANOVA with Bonferroni (B, G) or Tukey’s (C & E) correction for multiple comparisons, * P<0.05, ** P<0.005, **** P<0.0001.

To understand the dynamics of this phenotype we investigated the propensity of *lgc-41* mutant animals to leave a small food patch over time. The probability of worms to leave a food patch was measured at several time points after the animals were placed onto the assay plates, from 1 to 9 hours. At the timepoints 6 h and 9 h, we observed a significantly lower probability of leaving events for *lgc-41* mutant animals compared to N2 (Fig. 2D-E & S1A). Due to these results, only the 6 h timepoint was used in all further experiments.

To understand if the differences observed in the food leaving probability, or dispersal, could be due to defects in basal locomotion, the locomotion of *lgc-41* mutants was observed. In the presence of food, *lgc-41* mutants, displayed few significant differences in speed or body posture compared to N2 (Fig. S2). In addition, their ability to chemotax towards diacetyl, a known chemoattractant odour^18^, was evaluated and showed intact chemotaxis in the *lgc-41* (Fig. 2G).

A worm’s decision to leave a patch of food is dependent on the integration of sensory inputs into second order interneurons such as RIM^15^. To test whether LGC-41, may be required for this integration we measured the worm’s ability to detect the aversive odour 2-nonanone in the absence of food, as well as the food leaving behaviour in presence of the odour^19,20^. We saw no reduction in the ability of *lgc-41* mutants to detect 2-nonanone off food, instead they showed an increased aversion to 2-nonanone compared to N2 (Fig. 2G). Despite this increased aversion to 2-nonanone, *lgc-41* mutant worms took significantly longer to leave a food patch in the presence of 2-nonaone compared to N2 (Fig. 2I-L).

By reintroducing a wildtype copy of *lgc-41* under its own promoter we were able to significantly rescue all behavioural defects observed in the *lgc-41* mutant animals (Fig. 2B-L). Animals expressing the rescue construct under the *lgc-41* promoter had an increased dispersal rate in the absence of food, increased food leaving probability, and increased 2-nonanone induce food leaving rate. This strongly implicates LGC-41 in the control of food related behavioural states. To determine which specific neurons expressing *lgc-41* act to control these behaviours we next expressed a wildtype copy of *lgc-41* under the control of promoters expressed in ASI (*daf-7*), AQR/PQR (*gcy-37*), CEP (*dat-1*) and ASJ (*trx-1*). Interestingly, rescue of *lgc-41* in both ASI and AQR/PQR led to a small but significant increase in food leaving probability (Fig 2E) but did not rescue the dispersal (Fig. 2G, S1B) or 2-nonanone induced food leaving behaviours (Fig. 2I-L). This may be suggestive of a need for broad level expression of *lgc-41* in several neurons to control these complex behaviours.

### Betaine is Produced in *C. elegans* and its Synthesis Pathway Genes are Expressed in Neurons

Due to the broad expression of *lgc-41* within neurons and our observation of punctate LGC-41 protein expression in what appeared to be sensory ciliary endings, as well as in the densely synaptically connected nerve ring, we hypothesised that betaine may act upon LGC-41 in two contexts: either exogenously, for example via ingestion of betaine containing bacteria, or through endogenous production of betaine. We first sought to determine if wildtype worms are able to sense exogenous betaine. However, we saw no positive or negative chemotaxis towards betaine in wild-type or *lgc-41* mutant worms (Fig. S1C). Next, we sought to understand whether betaine may be produced within the worm and act as an endogenous ligand within the nervous system, by investigating the expression pattern of the putative betaine synthesis pathway genes *alh-9, alh-11* and *chdh-1* (Fig. 3A). The genes *alh-9, alh-11* were selected as putative betaine aldehyde dehydrogenase enzymes based upon their homology with ALDH7A1 (human betaine aldehyde dehydrogenase), with 65% and 27% identity respectively. *chdh-1* was selected as a putative choline dehydrogenase due to its 55% identity to CHDH (human choline dehydrogenase). Indeed, we identified expression of all 3 putative biosynthesis pathway genes in neurons (Fig. 3B). The RIM neurons expressed both *chdh-1* and *alh-11*, and interestingly the RIM neurons are presynaptic to the SAA neurons which we found to express the betaine-gated channel *lgc-41* (Fig. 1G), which could suggest a very direct role for betaine on SAA function.

**Figure 3.**
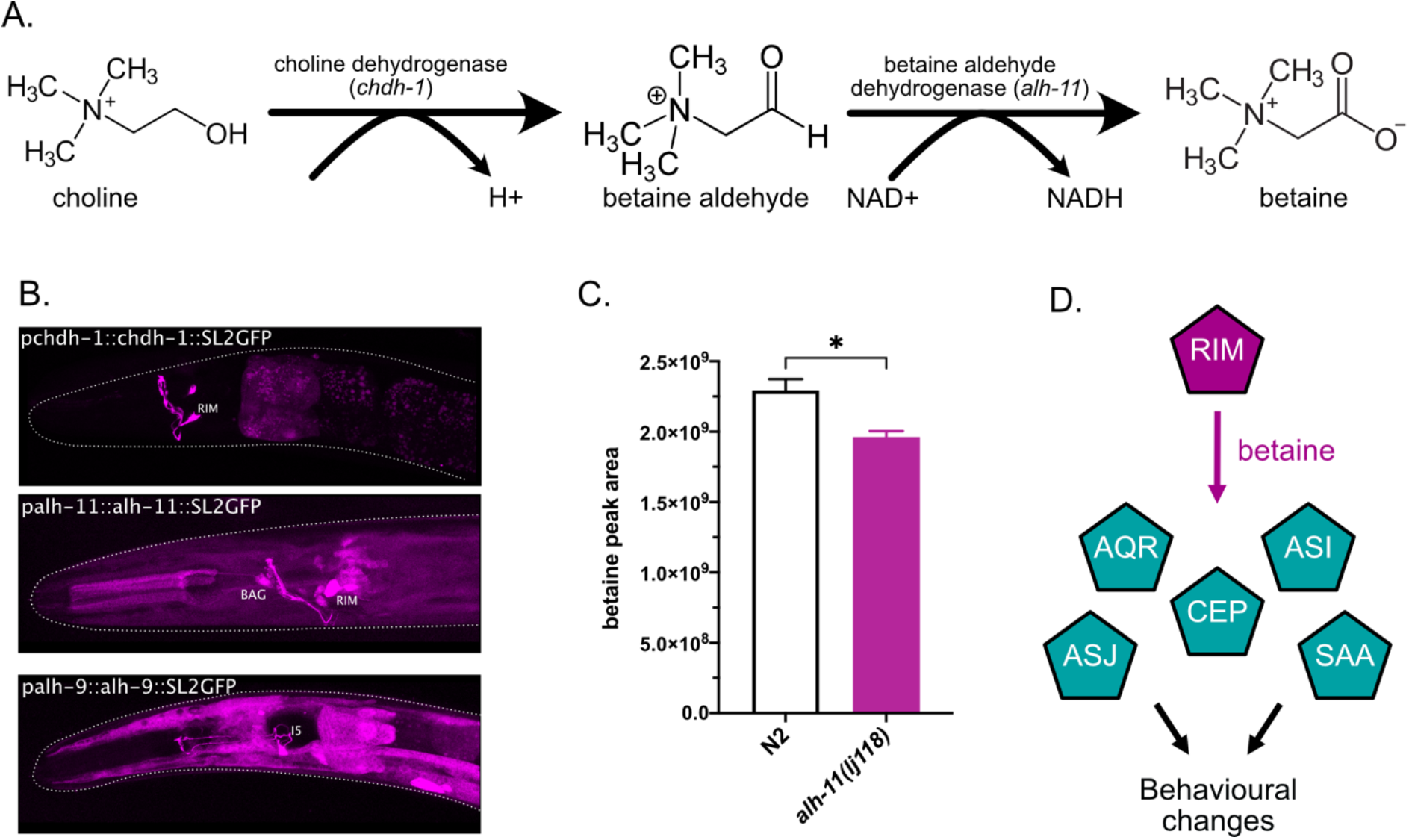
Neuronal *C. elegans* betaine synthesis pathway involves *alh-11*. (**A**) Schematic of betaine biosynthesis pathway and putative enzymes involved in each stage. (**B**) Fluorescent reporter of intercistonically spliced GFP driven under the promoter and genomic sequence of the putative betaine synthesis enzymes *alh-9, alh-11* and *chdh-1*. (**C**) Mass spectrometry analysis of betaine content in *C. elegans* wildtype or *alh-11* mutant preparation. The data shows peak area for betaine. Error bars represent SEM of 3 samples per genotype, *=P<0.05 calculated by unpaired t-test. (**D**) Schematic of betaine production and downstream neurons expressing LGC-41.

To further evaluate if one of the putative biosynthesis-pathway genes, *alh-11*, can influence the betaine levels found *in vivo* in *C. elegans*, we performed quantitative mass spectrometry analysis of N2 and *alh-11* mutant worms. In both samples high levels of betaine could be detected in the worm pellet, however, there was a significant reduction in betaine level in *alh-11* mutant worms compared to N2 (Fig, 3C). This strongly supports the notion that betaine is produced endogenously within the worm.

We next sought to understand if betaine may act *in vivo* to control the behaviours we observed in animals lacking the betaine receptor, LGC-41. To do this we characterised the behaviour of *alh-11* mutant animals, which have reduced endogenous betaine levels (Fig. 3C). Indeed, we observed strikingly similar behavioural defects in the *alh-11* mutants compared to *lgc-41* mutants. Like the animals lacking the betaine receptor, *alh-11* mutant animals had a reduced ability to disperse in the absence of food (Fig. 4A-C), and a reduced propensity to leave a food patch over time (Fig. 4D-E). Interestingly, however unlike the receptor mutants, *alh-11* mutants were not defective in 2-nonanone induced food leaving (Fig. 4I-K), suggesting other betaine sources may be relevant, for example, bacterial betaine production (Fig. S1E) or biosynthesis via *alh-9*, for which we were unable to generate null alleles due to homozygote lethality. Like *lgc-41* mutants, *alh-11* mutants also showed few significant differences in basal locomotion (Fig. S2) or chemotaxis, with the worms’ ability to chemotax away from the aversive odour, 2-nonanone, intact (Fig. 4F-H).

**Figure 4.**
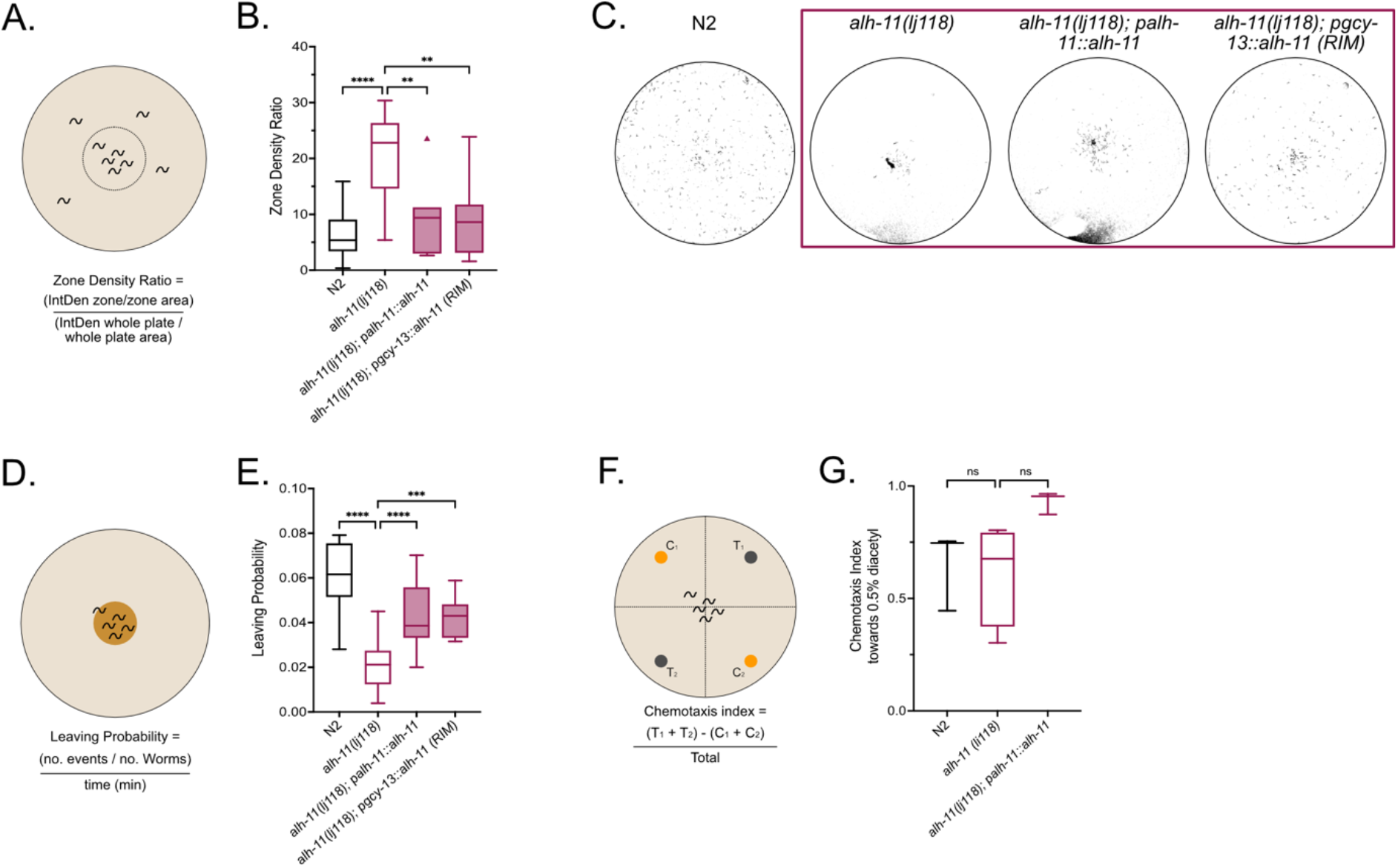
Endogenously produced betaine is required to promote global search and food leaving behaviours. Behavioural responses of N2, *alh-11(lj118), alh-11(lj118); palh-11::alh-11*, and *alh-11(lj118); pgcy-13(RIM)::alh-11* worms. (**A**) Schematic representation of the experimental design and (**B**) box plot of dispersal assay, in which central dispersal index is calculated and plotted. N=3-29 plates per genotype. (**C**) Representative images of threshold filtered plates used to calculated dispersal index. (**D**) Schematic representation of the experimental design and (**E**) box plot of food leaving probability 6h after being placed on the food patch. N=16-21 plates per genotype. (**F**) Schematic representation of experiment (**G**) Chemotaxis index towards 0.5% diacetyl. N= 3-5 plates per genotype. (**B, C, E, G**) Tukey’s blot plots and one-way ANOVA with Bonferroni (B, G) or Tukey’s (C & E) correction for multiple comparisons, ** P<0.005, *** P<0.001, **** P<0.0001.

Finally, due to our observation of punctate expression within synaptically dense regions, we sought to understand whether betaine may be produced and released from neurons, specifically the RIMs, which have been strongly implicated in food related behaviours and decision making^21^. We therefore reintroduced a wildtype copy of *alh-11* under the control of its own promoter and a RIM specific promoter (*gcy-13*) (Figure S1D). Expression under both promoters led to significant increases in dispersal rates (Fig 4B-C) and food leaving probabilities (Fig. 2E), which is the first *in vivo* evidence of a role for neuronally produced betaine as a neuromodulator. It also highlights a role for the RIM neurons in betaine transmission.

## Discussion

In this study we highlight a novel role for neuronal signalling using a non-classical neurotransmitter, betaine, and its receptor LGC-41. We demonstrate neuronal expression of putative betaine biosynthesis genes, and that the loss of one of these genes, *alh-11*, a putative betaine aldehyde dehydrogenase, caused a significant reduction in total betaine content in the body of *C. elegans*. We also showed that animals either lacking the betaine receptor, *lgc-41*, or the betaine synthesis gene, *alh-11*, displayed defects in multiple food search behaviours. These behavioural defects could be rescued by reintroducing the betaine synthesis gene, *alh-11*, specifically in the RIM interneurons, which have been implicated in the integration and decision-making processes during food search behaviour^21,22^, thereby strongly suggesting the need for neuronally produced betaine in the regulation of these behaviours. Interestingly, while *alh-11* defects could be rescued in a single neuron class, reintroduction of the receptor, *lgc-41*, in several other neuron classes did not result in strong behavioural rescues. The broad neuronal expression pattern along with a mixture of punctate and diffuse localisation of LGC-41 within neurons may suggest a complex interplay between synaptic and extra synaptic signalling via betaine and *lgc-41* which could likely vary between neuron classes and environmental states.

Previous work has implicated betaine in controlling normal locomotion through the muscular cationic betaine-gated channel ACR-23^9,10^. Interestingly, ACR-23 is distantly related to LGC-41, being more closely related to nAChRs. The presence of two independently evolved betaine-gated channels along with the presence of a betaine reuptake transporter, *snf-3*^9^, suggests a much broader role for betaine in the regulation of worm behaviours than previously recognised. The source of betaine acting via these two opposing receptors is not yet clear, our study suggests that neuronally produced betaine acts via LGC-41 in several neurons. However, worms lacking *alh-11*, the betaine synthesis enzyme, still contained betaine, and did not display any locomotion defects seen in *acr-23* mutant animals^9^, suggesting that the source of betaine acting upon *acr-23* in the muscle may be distinct from that acting on *lgc-41* in neurons. The remaining betaine present in the *alh-11* mutants may be contributed by the action of other enzymes and exogenous betaine synthesis by bacteria. Several types of bacteria, including *E. coli*, produce betaine to regulate their osmolarity^23^, and as such it is possible that betaine levels in bacteria may act as a marker of food quality for worms.

In mammals, betaine transport is closely associated with the transport of GABA^4^. In rodents, a lack of betaine has been shown to affect the balance of GABAergic transmission, causing to changes in memory formation^24^. Interestingly, there have also been reports of upregulation of betaine/GABA transporter expression in rodents after performing learning assays further indicating a specific role for betaine in the nervous system, separate from its osmoregulatory function. Many questions regarding the extent of betaine’s actions within animal nervous systems remain, for example whether betaine may be specifically loaded into and released from synaptic vesicles and the conditions and molecular mechanisms that may underlie such a process. However, our study strengthens the growing understanding that betaine can influence animal nervous systems far beyond its traditional roles and begins to piece together the neuronal mechanisms by which it can do so.

## Methods

### *C. elegans* culture

Unless otherwise specified, *C. elegans* worms were cultured on NGM agar plates with OP50^25^. A full list of strains used in this study can be found in Table S1. Unless otherwise specified 1-day adult worms were synchronised by placing L4 worms on culture plates overnight.

For metabolite extraction experiments the worms were cultured as follows: Ten *C. elegans* hermaphrodites were placed onto 6 cm NGM plates (each seeded with 200 *µ*L of OP50 E. coli grown to stationary phase in Lennox Broth) and incubated at 22 °C. After 96 hours, each plate was washed with 25 mL of S-complete medium into a 125 mL Erlenmeyer flask, 1 mL of OP50 E. coli suspension was added (*E. coli* cultures were grown to stationary phase in Terrific Broth, pelleted and resuspended at 1 g wet mass per 1 mL M9 buffer) and liquid cultures, shaking at 220 RPM 22 °C. After 70 hours, 6 mL bleach was added to each culture, along with 900 *µ*L 10 M NaOH and the mixture was shaken for 3 min to prepare eggs. Eggs were centrifuged at 1500x *g* for 30 seconds, supernatant discarded, and washed with 25 mL M9 buffer for three times. The pellet was re-suspended in final volume 5 mL M9 and transferred in a new 15 mL centrifuge tube. Eggs were placed on a rocker and allowed to hatch as L1 larvae for 24 hours at 22 °C. 100,000 L1s were counted and cultured in 25 mL cultures of S-complete with 1 mL of OP50 and incubated at 220 RPM 22 °C for 68 hours. Worms were then spun at 2000x *g* for 5 min and spent medium was separated from worm body pellet. Separated medium and worm pellet were flash frozen in liquid nitrogen and stored at -20 °C freezer until further processing.

### *Xenopus laevis* oocytes

Defolliculated oocytes from *Xenopus laevis* were obtained from EcoCyte Bioscience (Dortmund, Germany) and kept in ND96 (in mM: 96 NaCl, 1 MgCl_2_, 5 HEPES, 1.8 CaCl_2_, 2 KCl) solution at 16° C for 3-5 days.

### Molecular biology

The *lgc-41* cDNA sequence from wildtype N2 *C. elegans* were cloned from worm cDNA (generated by reverse transcription PCR from total worm RNA using Q5 polymerase (New England Biosciences)). Ion channel constructs for overexpression in *Xenopus* oocytes, were made from cDNA sequences and cloned into the KSM vector, the gene was inserted between *Xenopus* β-globin 5’ and 3’UTR regions and downstream of a T3 promoter using the HiFi assembly protocol (New England Biosciences). This protocol was also used for generating *C. elegans* expression constructs using the pDESTR4R3II vector. For fluorescent expression, *C. elegans* wildtype N2 gDNA was used for cloning gDNA sequences and the inclusion of GFP or mKate2 on the same plasmid after an intercistronic splice site (SL2 site). Unless otherwise specified promoter sequences consist of approximately 2kb of gDNA upstream of the start codon.

### CRISPR/CAS9-mediated gene manipulation

Endogenous tagging of the M3/4 cytosolic loop of *C. elegans* LGIC proteins with GFP was carried out either using the SapTrap protocol ^26,27^ for *lgc-41(lj119). C. elegans* genetic modifications of *lgc-41(lj115)* and *alh-11(lj118)* were made following ^28^.

### RNA synthesis and microinjection

The T3 mMessage mMachine transcription kit was used for *in vitro* synthesis of 5’ capped cRNA, the procedure was done according to manufacturer’s protocol (ThermoFischer Scientific). The cRNA was purified prior to injection using the GeneJET RNA purification kit (Thermo Fischer Scientific). *Xenopus* oocytes (Ecocyte) were individually placed into 96-well plates and injected with 50 nL of 500 ng/*µ*L RNA using the Roboinject system (Multi Channel Systems GmbH). A 1:1 ratio was used when two constructs were co-injected and with a remained total RNA concentration of 500 ng/*µ*L. The injected oocytes were incubated between 3-6 days post injection at 16°C in ND96 until the day of recording.

### Two-Electrode Voltage Clamp (TEVC)

recording and data analysis Using either the Robocyte2 system or a manual set up with an OC-725D amplifier (Multi Channel Systems GmbH), two-electrode voltage clamp recordings were carried out. Unless otherwise stated, oocytes were clamped at -60mV. Continuous recordings at 500Hz were taken during application of a panel of agonists or antagonists. Glass electrodes were pulled on a P1000 Micropipette Puller (Sutter), with a resistance ranging from 0.7-2 MΩ. Electrodes contained Ag|AgCl wires and backfilled with a 1.5M KCl and 1 M acetic mixture. Data was recorded using the RoboCyte2 control software, or with WinWCP for manual recordings, and filtered at 10 Hz.

Peak current for each dose was normalised to the oocyte maximum current using a custom-built python script ^29^. For dose response applications a protocol with 10s agonist application pulses with 60s of ND96 wash in between each dose was used. Using different batches of oocytes the data was gathered over at least two occasions. After normalisation the data was imported into Graphpad (Prism) and fitted to Hill equation (either a three or four parameter nonlinear) to obtain the highest degree of fit and to calculate the EC_50_. The EC_50_ dose of the primary agonist was used for dose response measurements as well as for ion selectivity recordings. Antagonist dose response protocols used 10s applications of agonist with antagonist present, followed by 60s of ND96 washes between doses. The agonist concentrations remained constant. IC_50_ values for each antagonist were calculated using a second custom built python script ^29^. After normalisation the data was fitted to a nonlinear Hill equation (either a three or four parameter) in Graphpad (Prism) to obtain the highest degree of fit and calculate the IC_50_.

A voltage ramp protocol from -80mV to +60mV (20mV/s) was used for ion selectivity measurments, in the presence of the primary agonist in three different solutions: ND96, NMDG (Na^+^ free) and Na Gluconate (low Cl-) solutions. The data was normalised to maximum current (I_max_) and the reversal potential (ΔE_Rev_) was calculated using a custom-built python script ^29^. Individual values, or mean, SD and n, for each construct was imported in GraphPad for further plotting and statistical analysis. Statistically significant differences in ΔE_Rev_ were calculated in GraphPad using a two-way ANOVA with Tukey’s correction for multiple comparisons. A representative normalised trace for each construct was also generated in Graphpad.

### Confocal and Cell ID

Worms were mounted onto 2% agarose in M9 pads and immobilised with 75 mM NaAz (in M9). A 63x water immersion lens on a Leica SP8 or STED or using a 40x oil immersion objective on a Zeiss LSM780 were used to acquire image stacks, these stacks were collapsed and max projections of z-stack images were generated in Fiji/Image J. Individial neurons were identified by cell shape, position or by crossing the reporter lines with the multicolour reference worm NeuroPAL ^30^.

### Metabolite Extraction

Worm samples frozen in liquid nitrogen were lyophilized to dryness. Dried worm pellets were homogenized by grinding using steel balls (2.5 mm balls, SPEX sample prep miniG 1600) at 1200 RPM for 3 minutes in 1-minute pulses and then extracted with 20 mL of methanol for 24 hours. The extracts were dried with a SpeedVac Vacuum Concentrator (ThermoFisher Scientific) and resuspended in 500 *µ*L MeOH, transferred to 1.7 mL Eppendorf tubes, centrifuged at 18,000 G for 10 minutes at 4 °C. Concentrated extracts were transferred to HPLC vials for HPLC-HRMS analysis.

### Mass Spectrometric Analysis

High resolution LC−MS analysis was performed on a ThermoFisher Scientific Vanquish Horizon UHPLC System coupled with a Thermo Q Exactive HF hybrid quadrupole-orbitrap high resolution mass spectrometer equipped with a HESI ion source. 1 *µ*L of extract was injected and separated using a water-acetonitrile gradient on an Agilent Zorbax Hilic Plus column (150 mm × 2.1 mm, particle size 1.8 *µ*m) maintained at 40 °C with a flow rate 0.3 mL/min. HPLC grade solvents were purchased from Fisher Scientific. Solvent A: 0.1% formic acid in water; Solvent B: 0.1% formic acid in acetonitrile. Analytes were separated using a gradient profile as follows: 2 min (95% B) → 20 min (50% B) → 22 min (10% B) → 25 min (10% B) → 27 min (95% B) →30 min (95% B). Mass spectrometer parameters: spray voltage 3.0 kV, capillary temperature 380 °C, prober heater temperature 300 °C; sheath, auxiliary, and spare gas 60, 20, and 2, respectively; S-lens RF level 50, resolution 240,000 at m/z 200, AGC target 3×106. The instrument was calibrated with positive and negative ion calibration solutions (ThermoFisher). Each sample was analyzed in positive and negative modes with m/z range 100 to 700. As a reference for betaine, betaine hydrochloride was purchased from Sigma-Aldrich. HRMS (ESI) m/z: calculated for C_5_H_12_NO_2_^+^, [M + H]^+^: 118.0863, found: 118.0860.

### Food Leaving Assays

Assays were carried out as described in ^31^. Briefly, 20 1-day adults were placed onto 55mm low-peptone agar assay plates seeded the day before with a 20*µ*L OP50 food patch in the centre. 15min videos were captured with DinoLite cameras (DinoLite, model no. AM7915MZTL) at the time points: 1h, 3h, 6h and 9h after placing worms onto the assay plates. Worm skeletons and trajectories were analysed using tierpsy ^32^, and the number of food leaving events was counted and expressed as food leaving probability. A leaving event was characterised by the full length of the worm’s body leaving the food patch. Food leaving probability was calculated as follows: Number of leaving events / the number of worms on the food patch at the start of the video / length of the video (minuets).

### Chemotaxis Assays

Assays were carried out as described in ^33^. Briefly, 1-day adults (synchronised by placing L4 worms on culture plates overnight, 4 days before the day of the assay) were washed 3 times in M9 and transferred onto 55mm CTX assay plates prepared with two test and two control substances, each mixed 1:1 with 1M NaAzide, totalling 2*µ*L per dot. 2-nonanone and diacetyl were diluted in 100% ethanol, betaine was dissolved in M9, in each case the diluent was used as the control substance. After 1h the number of worms in each quadrant was counted and the chemotaxis index calculated as follows: Chemotaxis Index = (No. of worms in Test 1 + No. of worms in Test 2) - (No. of worms in Test 1 + No. of worms in Test 2) / Total No. of worms. Worms that did not leave the centre of the plate were not counted, and only plates with a minimum of 50 worms were used in the analysis.

### 2-nonanone Induced Food Leaving Assays

Assays were carried out as described in ^20^. Briefly, 20 1-day adults were placed onto 55mm low peptone assay plates, seeded the previous day with 20*µ*L OP50. After 1 hour, and during video recording, a 1*µ*L dot of 100% 2-nonanone was placed directly next to, but not touching the food patch. The number of worms remaining on the food patch at 1min intervals was recorded for 10min, then again at 15min and calculated as a percentage of the total worms on the food before 2-nonanone addition.

### Dispersal Assays

1-day adult worms were synchronised by placing L4 worms on culture plates overnight, 4 days before the day of the assay. Worms were washed 4 times in M9 buffer to remove any bacteria and a densely packed 8*µ*L volume of worms was pipetted into the centre of the 55mm CTX agar assay plate. Once the M9 from the transfer dried, worms were left on the assay plates for 1h at which photos were taken of each plate. IntDen (integrated density) of a define centre zone, and the whole plate was calculated in Image J, after thresholding to remove background. Dispersal index was then calculated as follows: Dispersal Index = (IntDen zone/zone area) / (IntDen whole plate / whole plate area).

### Worm Tracking

Four 1-day adult worms were placed into the centre of the food patch on 55mm NGM plates seeded with 4*µ*L OP50 2 hours before. 15min videos were taken with DinoLite (DinoLite, model no. AM7915MZTL) cameras at 40X magnification and worm behaviour analysed with tierpsy ^32^. Where necessary trajectories for each worm were manually joined within the tierpsy GUI, the summary features for each trajectory were then exported and statistically significant differences were calculated on the 50^th^ percentile values for each feature using an unpaired t-test. Resulting p-values were plotted in a heatmap using the seaborn package in python, with every 4^th^ feature labelled.

### Data Analysis

Unless otherwise specified data was imported into GraphPad for further analysis and plotting. Unless otherwise specified, one-way ANOVA was used to calculate significant differences, using the Bonferroni (food leaving & dispersal) or Tukey’s (food leaving time course, chemotaxis) correction for multiple comparisons. *= P<0.05, **= P<0.005, ***=P<0.001, ****P<0.0001. Python scripts for TEVC analysis are deposited at GitHub at hiris25/TEVC-analysis-scripts. Data used for analysing TEVC experiments are available upon request from the Lead Contact.

## End notes

## Acknowledgements

We would like to acknowledge Lidia Ripoll Sanchez, Denise Walker and Amy Courtney, for assistance in cloning and the construction and maintenance of strains, as well as other past and present members of the Schafer lab for helpful discussions. This work was supported by grants from the Medical Research Council (MC-A023-5PB91), the Wellcome Trust (WT103784MA) and the National Institutes of Health (R35GM131877).

## Author Contributions

IH, JM, FCS and WRS designed experiments. IH, JM, JY and TSW performed experiments and analysed data. IH, JM & WRS wrote, and FCS, JY and TSW critically revised the manuscript.

## Additional Information

Supplementary Information is available for this paper.

## Supplementary Information

Table S1 – *C. elegans* strain list

## Supplementary Figure Titles

<u>Supplementary Figure 1:</u> ***lgc-41* and *alh-11* mutant worms show no significant differences in chemotaxis to betaine**.

<u>Supplementary Figure 2:</u> ***lgc-41* and *alh-11* mutant worms show few significant differences in basal locomotion**.

<u>Supplementary Figure 3:</u> ***lgc-41* but not *alh-11* mutant worms display a defect in 2-nonanone induced food leaving**.

