## Supplementary material for "Neuronally Produced Betaine Acts via a Novel Ligand Gated Ion Channel to Control Behavioural States": Table S1

Strain list

| Strain No. | Description |
|---|---|
| AQ4633 | alh-11(lj118) (bc 3x) |
| AQ4618 | lgc-41(lj115) (bc 4x) |
| AQ4658 | lgc-41(lj119) (bc 4x) |
| AQ4888 | liEx1479[palh-9::alh-9::SL2GFP; punc-122:RFP] on OH15262 |
| AQ4913 | ljEx1489[pchdh-1::chdh-1::SL2 GFP; punc-122::RFP] on OH15262 |
| AQ4550 | him-5(e1490); ljEx1305[palh-11(3kb)::alh-11::SL2-mkate2; punc-122::gfp] |
| AQ4525 | him-5(e1490); ljEx1297[palh-9(1.5kb)::alh-9::SL2-mKate2; punc-122::gfp] |
| AQ4523 | him-5(e1490); ljEx1295[pchdh-1(2kb)::chdh-1::SL2-mKate2; punc-122::gfp] |
| AQ4404 | him-5(e1490); ljEx1250[plgc-41(2kb)::lgc-41::SL2-mKate2; punc-122::gfp] |
| AQ4391 | ljEx1244[plgc-41(2kb)::mKate2::gpd-23'UTR; unc-122::gfp] |
| OH15262 | "NeuroPAL" otEx7057 |
| AQ5152 | alh-11(lj118); ljEx1595[palh-11(3kb)::alh-11::SL2GFP; punc-122::RFP] |
| AQ5177 | alh-11(lj118); ljEx1603[pgcy-13::alh-11::SL2GFP; punc-122::RFP] |
| AQ5135 | lgc-41(lj115), ljEx1584[pgcy-37::lgc-41::SL2mKate; punc-122::gfp] |
| AQ5088 | lgc-41(lj115), ljEx1576[pdaf7(4.5kb)::lgc-41::Sl2mKate; punc-122::gfp] |
| AQ5224 | lgc-41(lj115); ljEx1619[pdat-1::lgc-41::Sl2mKate; punc-122::gfp] |
| AQ5225 | lgc-41(lj115); ljEx1619[ptrx-1::lgc-41::Sl2mKate; punc-122::gfp] |

Note:
*chdh-1* previously called *C34C6.4*
