## Supplemental Figures for "Neuronally Produced Betaine Acts via a Novel Ligand Gated Ion Channel to Control Behavioural States"

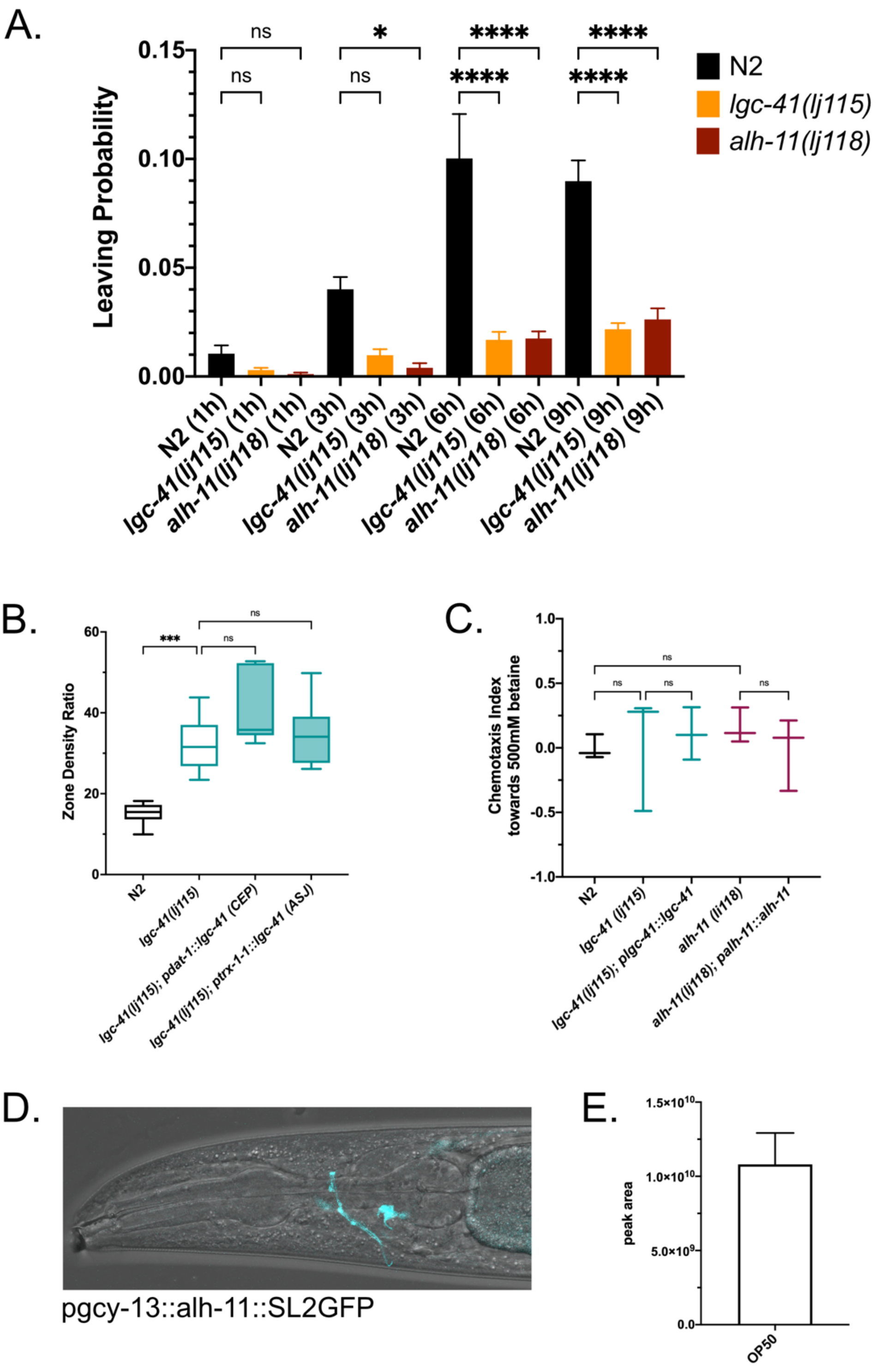

Supplementary Figure 1. ***lgc-41* and *alh-11* mutant worms show no significant differences in chemotaxis to betaine.** (**A**) Food leaving probability time course, n=5 plates for each genotype and each time point. Significance calculated by 2way ANOVA with Tukey's correction for multiple comparisons. * P<0.05, **** P<0.0001. (**B**) Box plot of dispersal propensity of N2, *lgc-41(lj115), pdat-1(CEP)::lgc-41* and *lgc-41(lj115); ptrx-1(ASJ)::lgc-41* worms, in which central zone density is calculated and plotted. N=6-8 plates per genotype. Tukey's blot plots and one-way ANOVA with Tukey correction for multiple comparisons, ***=P<0.001. (**C**) Chemotaxis towards or away from 500 mM betaine in wildtype (N2) or *lgc-41(lj115)*, *alh-11(lj118)* mutants or genomic rescue lines. n=3-5 per genotype, significance calculated by one way ANOVA with Tukey's correction for multiple comparisons. (**D**) Fluorescent reporter image of RIM specific expression of *alh-11* under the control of *pgcy-13*. (**E**) Mass spectrometry analysis of betaine content in *E.coli (*OP50). The data shows peak area for betaine. Error bars represent SEM of 5 samples.

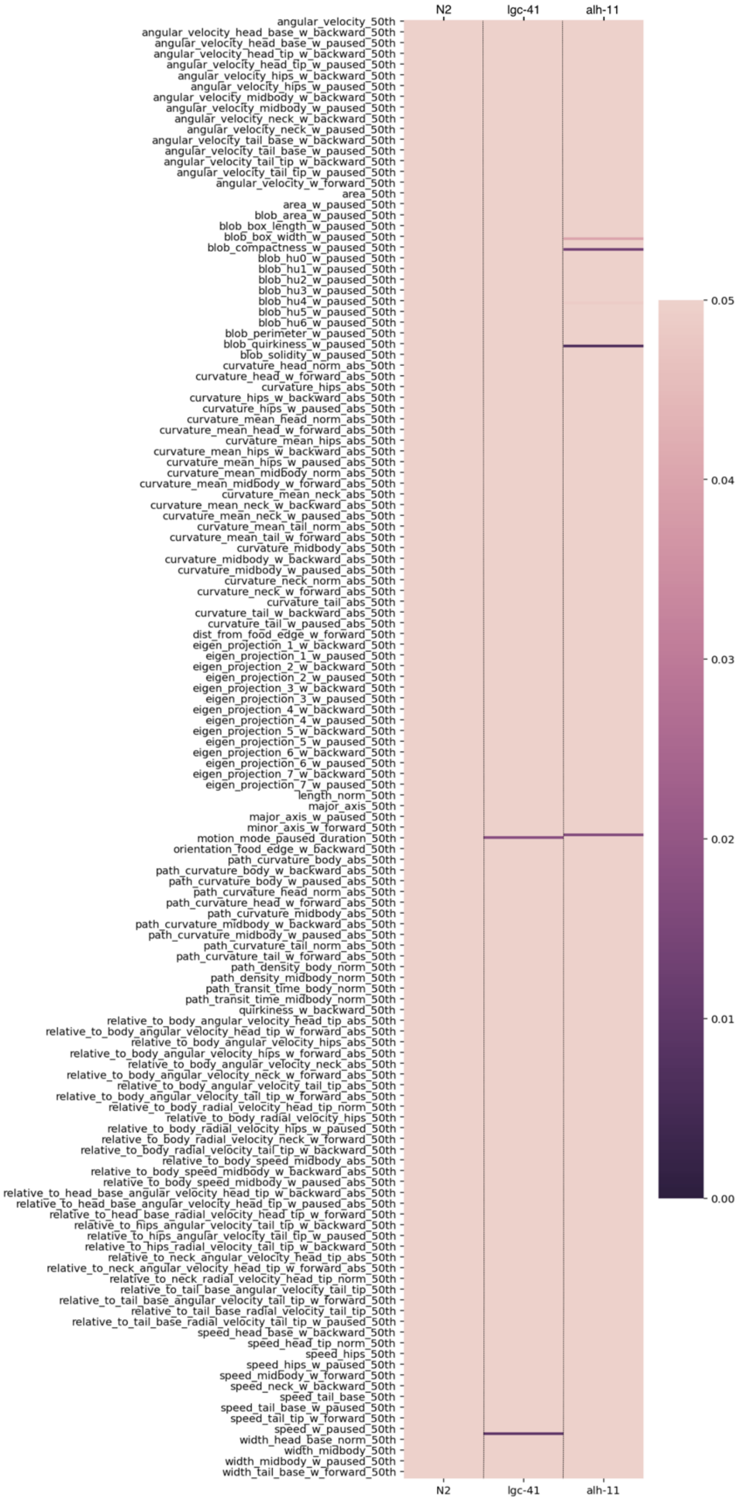


<u>Supplementary Figure 2.</u> ***lgc-41* and *alh-11* mutant worms show few significant differences in basal locomotion.** Heat map showing significance values of locomotion

tracking features of *lgc-41(lj115)* and *alh-11(lj118)* compared to wildtype (N2) worms in the presence of food. N=6 plates per genotype.

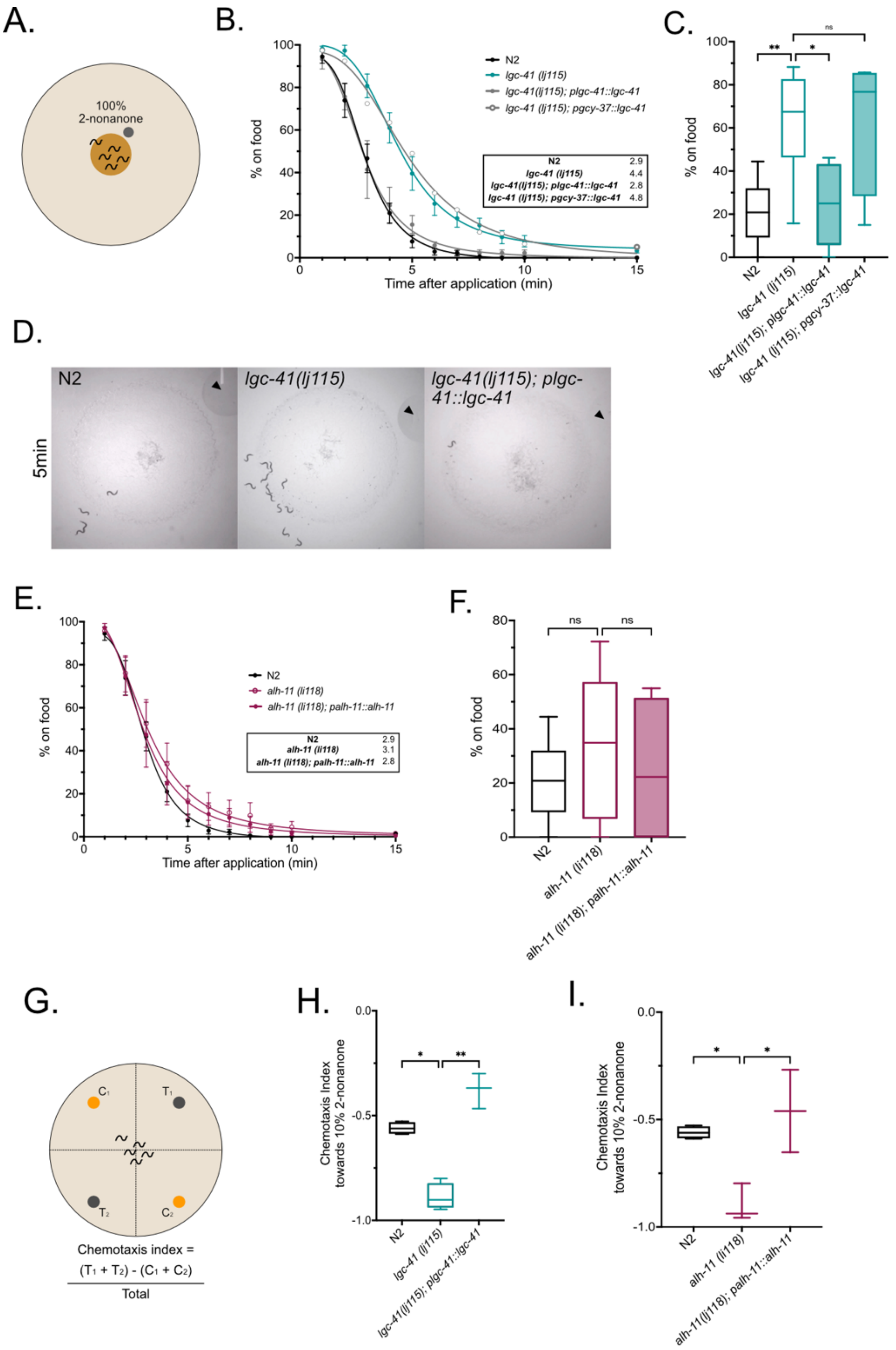


<u>Supplementary Figure 3.</u> ***lgc-41* but not *alh-11* mutant worms display a defect in 2-nonanone induced food leaving.** Behavioural responses of N2, *lgc-41(lj115)*, *lgc-41(lj115); plgc-41::lgc-41*, *lgc-41(lj115); pgcy-37(AQR/PQR)::lgc-41*, *alh-11(lj118)* and *alh-11(lj118);*

*palh-11::alh-11* worms. (**A**) Schematic representation of the experimental design of 100% 2-nonanone induced food leaving, the graph (**B, E**) displays percentage of worms on food each minuet after 2-nonanone application for 15min, curves fit with a four-parameter variable slope, insert gives time in minuets when 50% of worms have left the food patch. N=5-12 plates per genotype, error bars represent SEM. (**C, F**) Box plot of percentage of worms on the food patch 4min after 2-nonanone application. N=5-12 plates per genotype. (**D**) Representative images of N2, *lgc-41* mutant and *lgc-41* rescue worms 5min after 2-nonanone application, black arrow indicated 2-nonanone droplet. (**G**) Schematic representation of experiment (**H, I**) box plot of chemotaxis index towards 10% 2-nonanone. N=3-5 plates per genotype. (**C, F, H, I**) Tukey's blot plots and one-way ANOVA with Bonferroni (C, F) or Tukey's (H, I) correction for multiple comparisons, * $P<0.05$, ** $P<0.005$.
